# Fatal Leptospirosis in Southern Sea Otters from Central California: Pathologic Findings and Detection of *Leptospira interrogans*

**DOI:** 10.64898/2026.01.08.698541

**Authors:** Margaret E. Martinez, Pádraig J. Duignan, Katherine C. Prager, Cara L. Field, Mary E. Gomes, Rinosh Mani, Lilian Carswell, Tim Tinker, Joseph Tomoleoni, Michael Murray, Ri K. Chang, James O. Lloyd-Smith, Melissa Miller

**Affiliations:** The Marine Mammal Center, 2000 Bunker Road, Sausalito CA 94960 USA; Marine Wildlife Veterinary Care and Research Center, California Department of Fish and Wildlife.151 McAllister Way, Santa Cruz, CA, 95060 USA; Department of Ecology and Evolutionary Biology, University of California, Los Angeles, 610 Charles E Young Drive East, Los Angeles, CA 90095 USA; Monterey Bay Aquarium, 886 Cannery Row, Monterey, CA 93940 USA; Veterinary Diagnostic Laboratory, Michigan State University 4125 Beaumont Rd, Lansing, MI 48910 USA; Wildlife Health Center, UC Davis School of Veterinary Medicine, 1089 Veterinary Medicine Drive, Davis CA, 95616 USA; U. S. Fish and Wildlife Service, Ventura Fish and Wildlife Office, 2493 Portola Road, Suite B, Ventura, CA, 93003-7726 USA; Nhydra Ecological Consulting, 11 Parklea Dr, Head of St Margaret’s Bay, NS, B3Z2G6, Canada; U. S. Geological Survey, Western Ecological Research Center, 2885 Mission St., Santa Cruz, CA 95060 USA

**Keywords:** Southern sea otter (Enhydra lutris), leptospirosis, Leptospira interrogans serovar Pomona, Kidney Diseases, Liver Diseases, Marine mammals, Wildlife animals

## Abstract

*Leptospira interrogans* serovar Pomona infections cause periodic outbreaks among California sea lions (CSL; *Zalophus californianus*) and sporadic deaths in phocids; however, frequency of infection and associated health impacts are uncharacterized among sympatric threatened southern sea otters (SSO; *Enhydra lutris nereis*), which are important sentinels of coastal health. Given the broad impacts of *L. interrogans* on other marine mammals, our objective was to screen selected SSO for infection, determine whether leptospirosis contributes to SSO mortality, and describe leptospiral-associated lesions. A retrospective review (2005-2025) yielded 19 candidates that underwent case review, including *Leptospira* immunohistochemistry (IHC), serology, and polymerase chain reaction (PCR), with a special focus on renal and hepatic lesions. Kidney was PCR-positive for 74% (14/19) of suspected cases and for eight of these, DNA sequence-based serogroup typing detected *L. interrogans* serogroup Pomona. Seven of the 14 PCR-positive leptospirosis cases were classed as fatal based on positive renal IHC and moderate to severe tubulointerstitial nephritis. All fatal cases had anti-*L. interrogans* serovar Pomona antibody titers ≥1:25,600. All remaining (n=7) PCR-positive cases were considered nonfatal leptospirosis cases due to minimal and/or unrelated renal lesions, and negative IHC. Nonfatal *Leptospira-*infected cases ranged from seronegative to low positive (1:400) for serovar Pomona. Antibody titers for *Leptospira* PCR-negative cases were negative. For fatal cases, gross renal changes were often inapparent or characterized by miliary white cortical foci. Renal histologic lesions included tubulointerstitial nephritis, acute tubular necrosis and suppurative tubulitis with intratubular bacteria in fatal cases, and positive IHC staining for leptospiral antigen in lesions. Gross hepatic changes were also inapparent in fatal cases and histologic lesions were rare, characterized in one animal by hepatocellular disassociation and in two sea otters by limited leptospiral antigen detection by IHC. Most *Leptospira*-infected sea otters (71%, 10/14) stranded during higher rainfall months in California, suggesting possible land-to-sea transmission from terrestrial hosts. Given these findings, and because *L. interrogans* serovar Pomona infections have been confirmed in sympatric CSLs and terrestrial mammals from adjacent watersheds, focused investigation of potential marine and terrestrial disease transmission dynamics could provide new information to reduce mortalities of SSO.

## 1 Introduction

Southern sea otters (SSO, *Enhydra lutris nereis*), federally listed as threatened in the United States, are a keystone sub-species in the California Current ecosystem and are recognized sentinels of ocean health (Kreuder et al., 2003; Miller et al., 2020b). These otters reside in geographically localized areas near the coastline and occupy relatively small home ranges, and they are often at risk from coastal anthropogenic development and land-to-sea pathogen transmission (Breed et al., 2017; Burgess et al., 2020; Kreuder et al., 2003; Miller et al., 2002; Shapiro et al., 2012, 2010; Tarjan and Tinker, 2016). Several comprehensive studies have characterized the prevalence of various land- and marine-origin diseases and anthropogenic stressors that can impede SSO population recovery (Burgess et al., 2020; Kreuder et al., 2003; Miller et al., 2020b, 2010; Shapiro et al., 2019). One study assessed serological evidence of exposure to the bacterial pathogen *Leptospira* spp. and reported seroprevalence of 10% in stranded SSO and 1.6% in free-ranging animals sampled from 1995-2000 (Hanni et al., 2003); however, infection with *Leptospira* and any consequent disease remain poorly characterized in this subspecies.

Leptospirosis, a bacterial disease caused by infection with spirochetes in the genus *Leptospira*, affects many mammalian species worldwide, including humans, and often presents as acute renal failure, hepatopathy, hemodynamic disturbances as well as other pathologic changes. The lesion pattern can vary widely in relation to the mammalian host and bacterial serovar (Chacko et al., 2021; Haake and Levett, 2015; Rajapakse, 2022). The kidney is most often targeted, with the classic histologic lesion, tubulointerstitial nephritis, often most concentrated in the renal cortex and at the corticomedullary junction (Maxie, 2016). However, *Leptospira* spp. infections can exhibit a wide spectrum of severity and may present with few or mild signs or even no clinical disease or hematological abnormalities (Chacko et al., 2021; Haake & Levett, 2015; Prager et al., 2013, 2020). Transmission of *Leptospira* is predominantly via urinary shedding and subsequent exposure via mucous membranes or broken skin (Chacko et al., 2021; Gostic et al., 2019; Haake and Levett, 2015). Pathogenic leptospires can remain infectious in fresh water or soil for weeks to months, but survival in salt water is much shorter (Bierque et al., 2020; Cilia et al., 2020).

California sea lions (CSL, *Zalophus californianus*) residing along the eastern Pacific coast are a well-documented host of *L. interrogans* serovar Pomona and have seasonal outbreaks each year and a historical pattern of major epizootics every 3-5 years (Gulland et al., 1996; Lloyd-Smith et al., 2007). Infected CSLs exhibit a spectrum of clinical manifestations, and the outcome can range from no clinical disease to fatality (Prager et al., 2020; Whitmer et al., 2021a). The highest incidence of fatal leptospirosis occurs in juvenile CSLs, with a bias toward males, and documented strandings due to leptospirosis are concentrated in central California (Greig, et al., 2005; Gulland et al., 1996). Horizontal transmission via urine is postulated to occur at haulout sites or in brackish waters, and population-scale persistence of the pathogen is likely facilitated by those without clinical disease that shed leptospires (Buhnerkempe et al., 2017; Prager et al., 2013, 2020).

Several recent serologic and disease studies have also documented the circulation of pathogenic *Leptospira* in coastal terrestrial mammals in California. High prevalence of infection has been reported in mesocarnivores and squirrels, with elevated levels in the central coast and indications that the agent was *L. interrogans* serovar Pomona (Straub et al., 2020; Straub and Foley, 2020). A survey in coastal southern California confirmed widespread infection with *L. interrogans* serovar Pomona in raccoons, coyotes and striped skunks, with genotypes closely related to those detected in CSLs, suggesting possible land-to-sea transmission (Helman et al., 2023).

SSOs thus have two potential sources of exposure to pathogenic *Leptospira*, via habitat sharing with infected CSLs or land-to-sea transmission from abundant terrestrial reservoirs. Yet *Leptospira* spp. infection and mortality have not been investigated in SSO. A recent report of leptospirosis in six northern sea otters (NSO, *Enhydra lutris kenyoni*) in Washington, USA, described tubulointerstitial nephritis, positive immunolabeling of kidney for leptospiral organisms, and seropositivity for *L. interrogans* serovar Pomona (Knowles et al., 2020). Given the broad impacts of *L. interrogans* infection on sympatric CSL and coastal terrestrial mammals, our objectives were to screen a selection of SSO cases with suspected *Leptospira* infection, determine whether it contributed to mortality, and describe *Leptospira*-associated disease.

## 2 Material and Methods

### 2.1 Study Animals and Ethics Statement

This study included: a) SSO that stranded and died naturally or were subsequently humanely euthanized due to poor prognosis by Marine Wildlife Veterinary Care and Research Center (MWVCRC; Santa Cruz, CA), The Marine Mammal Center (TMMC; Sausalito, CA), Monterey Bay Aquarium (MBA; Monterey, CA), or other authorized members of the Southern Sea Otter Stranding Network (SSOSN); b) SSO carcasses that were found dead along the central Californian coast and recovered by CDFW, TMMC, MBA, or other SSOSN staff.

The recovery of carcasses, humane euthanasia of SSO, and sample collection by MWVCRC, TMMC, and MBA were performed in accordance with Section 109(h) of the U.S. Marine Mammal Protection Act (MMPA) and the U.S. Fish and Wildlife Service (USFWS) regulations implementing the MMPA at 50 CFR 18.22(a), and in accordance with the Service’s regulations implementing the U.S. Endangered Species Act at 50 CFR 17.21(c)(3). Permission for animal or carcass recovery, rehabilitation, humane euthanasia, and sample collection was granted by USFWS permits MA101713-1 and PER4324646 for TMMC, and MA032027 and MA186914 for MBA. No Institutional Animal Care and Use Committee protocol was needed as samples were collected from stranded, hospitalized SSO during their routine care Animals were sedated or anesthetized with a combination of fentanyl citrate (0.22-0.33 mg/kg IM) and midazolam HCl (0.07-0.11 mg/kg IM) which was reversed with naltrexone three to five times the fentanyl dose. Supplemental oxygen was provided either via face mask or tracheal intubation. After otters were anesthetized, they were euthanized in accordance with the current American Veterinary Medical Association guideline (AVMA, 2020). After being sedated with fentanyl/midazolam, they were euthanized with intravenous or intra-cardiac injection of a euthanasia solution containing pentobarbital doses recommended by the manufacturer.

### 2.2 Case Selection

A retrospective search of MWVCRC’s and TMMC’s databases was conducted to identify SSO submissions between 2005-2025 that had clinical signs or laboratory or pathologic findings suggestive of leptospirosis. Case inclusion criteria included animals that were necropsied with only minimal or moderate decomposition, had renal and hepatic tissues available for histologic review, cryopreserved kidney available for diagnostic screening, and disease processes that included one or more of the following key words: nephropathy, nephritis, interstitial nephritis, tubulointerstitial nephritis, hepatopathy, hepatitis, leptospirosis, *Leptospira* and icterus. Based on the inclusion criteria, 19 cases were selected for additional leptospiral testing to categorize the leptospiral status (see case definitions section). The case selection was intentionally biased for increased chances of identifying leptospirosis cases in this population to serve as a pilot study and guidance for future epidemiologic studies. Age class categories (immature [6 months-1 year], subadult [1–3 years], adult [4+ years]) were determined based on dentition as previously described (Miller et al., 2020b).

### 2.3 Case Definitions

For each animal fitting the above criteria, all archival case material, including clinical summaries, necropsy reports, histologic slides, histologic reports, final pathology reports, clinical, necropsy, histologic and ultrasonographic images, and sample archives were reviewed by a veterinary pathologist. Additional diagnostics, such as *Leptospira* PCR, immunohistochemistry (IHC), and serology were completed and were reviewed in context with the case material. Particular attention was given to renal and hepatic lesions. Major causes of death/stranding were determined by the pathologist after complete case material review and on lesion severity. Leptospirosis was not determined to be a major contributor to death/stranding when renal or hepatic changes were consistent with incidental findings (e.g. mild interstitial nephritis) or secondary to other major causes of death/stranding (e.g. bile cast nephropathy secondary to microcystin toxicosis).

Following case review, enrolled cases were classified into three groups: (1) Fatal leptospirosis cases had PCR-positive kidneys, moderate to severe tubulointerstitial nephritis, intralesional immunopositive leptospiral organisms, and evidence that leptospiral related disease was a major cause of death/stranding. (2) Nonfatal leptospiral cases had PCR-positive kidneys, negative IHC, and no evidence of leptospiral related lesions as a major cause of death/stranding. (3) Negative cases had PCR-negative kidneys, negative IHC, and no evidence of leptospiral related lesions as a major cause of death/stranding.

### 2.4 Gross, Histologic, and *Leptospira* Immunohistochemical Findings

SSOs that stranded throughout central California, USA or died under human care were necropsied at MWVCRC or TMMC using standardized protocols (Miller et al., 2020b). All major organs were sampled, formalin fixed and processed for histologic examination as previously described (Kreuder et al., 2003). Renal changes were scored on a 0-3 scale with 0 equating to none, 1 equating with mild and/or <25% of tissue affected, 2 equating with moderate and/or 25-50% of tissue affected, and 3 equating with severe and/or >50% of tissue affected for each of the following lesions: interstitial only nephritis, tubulointerstitial nephritis, tubulitis, acute tubular necrosis/attenuated tubular epithelium, tubular casts, other tubular changes, and glomerular or vascular changes. Changes were scored for severity within the renal cortex, medullae and papillae. Overall renal histologic morphologic diagnoses were also assigned. Specific tubular and glomerular or vascular changes were noted as well as the cell type(s) present when scoring each change. Liver was also examined microscopically and was assigned an overall histologic morphologic diagnosis.

IHC was performed at the California Animal Health and Food Safety Laboratory (CAHFS, Davis, CA, USA) on paraffin embedded, formalin fixed liver and kidney for all cases. The primary antibody was a *Leptospira* antibody (Rabbit origin multivalent fluorescent antibody conjugate, FITC-bound, LEP-FAC, National Veterinary Services Laboratories, Ames, IA, 50010, USA) with a dilution factor of 1:3000 and no antigen retrieval. After primary incubation for 45 min, excess primary antibody was rinsed off with wash buffer then the secondary antibody was applied (Dako EnVision+ HRP Anti-Rabbit K4003, Agilent, Santa Clara, CA 95051, USA) and incubated for 30 min. After two further washes, chromogen stain (BioCare Romulin AEC, 810, Pacheco, CA 94553, USA) was applied. A final wash was conducted to remove the chromogen and diluted Gil Hematoxylin was applied as a counterstain. Finally, renal slides and controls were evaluated for presence or absence of granular staining and scored as 0 equating to no staining, 1 for <25% tissue staining, 2 equating to 25-50% tissue staining, and 3 for >50% tissue staining. Staining was scored and evaluated for the cortices, medullae and papillae, and a specified location of staining recorded for the interstitial inflammatory cells, tubular epithelium, or tubular luminal contents. The liver was evaluated based on the location and intensity of positive immunohistochemical staining.

### 2.5 Imaging and Clinical pathology

In a subset of live-stranded animals presenting for clinical care, anesthesia was conducted with intramuscular fentanyl (0.22 mg/kg, IM, Central Avenue Pharmacy, Pacific Grove, CA 93950 USA) and midazolam (0.07 mg/kg, IM, Hikma Pharmaceuticals, Barkeley Heights, NJ 07922 USA) following a standard protocol at Monterey Bay Aquarium (MBA) (Monson et al., 2001) or midazolam (0.2mg/kg, IM, Pfizer, Lake Forest, IL 60045 USA), butorphanol (0.2mg/kg, IM, Zoetis Inc, Kalamazoo, MI 49007 USA) and dexmedetomidine (0.03mg/kg, IM, Zoetis Inc) at TMMC. Abdominal ultrasonography (U/S) was performed using an HM70 EVO portable ultrasound and microconvex 4-10 MHz transducer (Samsung, Ridgefield Park, NJ 07660 USA).

Whole blood was collected via the jugular vein under anesthesia or immediately prior to euthanasia using a 19-gauge, 19 mm butterfly needle connected to vacutainer tubes (Becton Dickinson, Oakville, ON LH6 6R5, Canada). Blood tubes containing ethylenediaminetetraacetic acid (K2 EDTA) were used for complete blood counts (CBC), and tubes containing no additive were collected for serum biochemistry. Coagulated whole blood was centrifuged at 800xg for 10 min to generate serum, which was either analyzed within four hours, or cryopreserved at −80°C for later analysis. Serum biochemistry was performed using an Axcel clinical chemistry analyzer (Alfa Wasserman-West, Caldwell, NJ 07006, USA) at TMMC or was submitted to IDEXX (Westbrook, ME 04092 USA). Complete blood counts were analyzed by a Vet ABC Plus analyzer (SCIL Vet America, Gurnee, IL 60031, USA) at TMMC or submitted to IDEXX. White blood cell differential counts and cell morphology were read manually from blood smears stained with Wright-Giemsa. Results were compared to published ranges derived from healthy free-ranging SSO (Williams and Pulley, 1983).

### 2.6 *Leptospira* Serology

Antemortem or postmortem serum from the nearest timepoint to euthanasia was submitted for serology; where possible, antemortem serum testing was prioritized to minimize effects of postmortem hemolysis. Microscopic agglutination testing (MAT) was performed at CAHFS (n=6) or at the Centers for Disease Control and Prevention (CDC), Atlanta, Georgia (n=8). Results were compiled for the case series as results are qualitatively the same and typically within a couple dilutions of each other (Mummah et al., 2024). Both laboratories participate in the international proficiency testing to ensure results are consistent among laboratories. Reference strains of live-cultured *Leptospira* spp. were used to assess serum anti-*Leptospira* antibody titers. CAHFS ran samples against a six serovar *Leptospira* panel, whereas the CDC ran samples against 19 serovars. MAT protocol details are as previously published (Prager et al., 2013, 2015, 2020) and are briefly summarized below.

Endpoint titers were determined by starting with an initial serum dilution of 1:100 followed by two-fold serial dilutions until at least 50% of the cells for the strain were agglutinated (Prager et al., 2013). Titers of 1:100 or higher were considered seropositive. The 19-serovar panel (CDC) included: Australis, Autumnalis, Ballum, Bataviae, Bratislava, Borincana, Canicola, Celledoni, Cynopteri, Djasiman, Grippotyphosa, Icterohaemorrhagiae, Javanica, Georgia, Mankarso, Pomona, Pyrogenes, Tarassovi and Wolffi. The six serovar panel (CAHFS) included: Bratislava, Canicola, Gryppotyphosa, Hardjo, Icterohaemorrhagiae and Pomona.

### 2.7 *Leptospira* PCR and typing

At necropsy, kidney and liver were collected in whirl packs or cryovials and stored at −80°C until shipping frozen to Michigan State University College of Veterinary Medicine (Lansing, MI, 48824, USA). DNA was extracted from kidney cryopreserved at −80C using Qiagen DNeasy Blood and Tissue kit. Screening for Leptospira DNA was performed using a real time PCR targeting the LipL 32 gene of pathogenic Leptospira spp. as previously described (Miotto et al., 2018; Rojas et al., 2010). As per Rojas et al. the limit of detection is 3 genome copies per reaction. In brief, 2uL template DNA was added to 10uL 2x Taqman universal PCR master mix and 0.8uM (final concentration) forward and reverse primer (lipl32-F, 5’-TAAAGCCAGGACAAGCGCC-3’; lipl32-R, 5’- CGCCTGGYTCMCCGATT-3’) and 0.4uM probe (lipl32-P, FAM/ATTTCCCCAACAGGCG/MGBNFQ) and DNAase free water to make up to a 20uL PCR reaction. The real-time PCR was performed on ABI 7500 (95°C for 10 minutes followed by 45 cycles of 95C for 0.15s and 60C for 1min). Positive samples were further analyzed using a serogroup-specific PCR assay. The serogroup-specific PCR amplified the LigB C-terminus gene of Leptospira. The resulting PCR product was Sanger sequenced for identification (Saraullo et al., 2021; Stoddard et al., 2009; Wells et al., 2024).

### 2.8 Microcystin

Microcystin (toxins produced by cyanobacteria) testing was previously performed on several cases with hepatic lesions suggestive of toxicosis or leptospirosis. High performance liquid chromatography tandem mass spectrometry was used to analyze gastrointestinal content, feces and/or urine for the presence of microcystin at either the California Department of Fish and Wildlife Water Pollution Control Laboratory, or at the University of California, Santa Cruz, using previously published protocols (Miller et al., 2010).

## 3 Results

### 3.1 Case selection

Nineteen SSO (16 from MWVCRC, and three from TMMC) necropsied from 2005 through 2025 met all criteria for case inclusion. This sample population consisted of 13 males and six females, with age classes ranging from subadult through aged adult (Table 1). Nearly all SSO (95%; 18/19) were fresh at the time of necropsy; one (case 11) was moderately decomposed. Eight animals stranded alive were euthanized, one stranded alive but died prior to arrival at the rehabilitation center, and the remainder were found dead. Strandings occurred in all calendar months except July and December.

**Table 1.** Demography and selected diagnostic test results for necropsied, *Leptospira-* infected and uninfected southern sea otters. MAT = Microagglutination test. CBD = Cannot be determined (due to hemolyzed serum sample). Asterisk = DNA sequence-based serogroup typing determined *L. interrogans* serogroup Pomona.

| Case | *Leptospira*<br>Case<br>Definition | Age Class | Sex | Stranding<br>Date | Case<br>History | Kidney<br>*Leptospira*<br>PCR Result | *L. interrogans*<br>serovar Pomona<br>MAT titer |
| --- | --- | --- | --- | --- | --- | --- | --- |
| 1 | Fatal | Aged<br>Adult | Male | 5/15/2011 | Euthanized | Positive* | 51,200 |
| 2 | Fatal | Adult | Female | 10/24/2011 | Euthanized | Positive* | 25,600 |
| 3 | Fatal | Adult | Male | 4/23/2015 | Euthanized | Positive* | 51,200 |
| 4 | Fatal | Subadult | Female | 9/14/2023 | Euthanized | Positive | 102,400 |
| 5 | Fatal | Subadult | Male | 1/27/2025 | Found<br>Dead | Positive* | 102,401 |
| 6 | Fatal | Adult | Male | 2/2/2025 | Dead on<br>Arrival | Positive* | 102,402 |
| 7 | Fatal | Adult | Male | 5/16/2025 | Euthanized | Positive* | 51,200 |
| 8 | Nonfatal | Subadult | Male | 1/30/2005 | Found dead | Positive | CBD |
| 9 | Nonfatal | Adult | Male | 1/30/2007 | Found dead | Positive | CBD |
| 10 | Nonfatal | Adult | Male | 6/9/2007 | Euthanized | Positive | CBD |
| 11 | Nonfatal | Aged<br>Adult | Male | 4/10/2010 | Found dead | Positive | CBD |
| 12 | Nonfatal | Aged Adult | Male | 4/20/2010 | Found dead | Positive* | 0 |
| 13 | Nonfatal | Adult | Female | 3/24/2014 | Found dead | Positive | 400 |
| 14 | Nonfatal | Adult | Female | 1/5/2017 | Found dead | Positive* | 0 |
| 15 | Negative | Aged Adult | Male | 6/13/2005 | Found dead | Negative | 0 |
| 16 | Negative | Adult | Male | 2/22/2006 | Found dead | Negative | 0 |
| 17 | Negative | Adult | Male | 8/30/2007 | Found dead | Negative | CBD |
| 18 | Negative | Subadult to Adult | Female | 2/26/2023 | Euthanized | Negative | 0 |
| 19 | Negative | Aged Adult | Female | 8/14/2023 | Euthanized | Negative | 0 |

### 3.2 Case Definitions and Demographics

PCR screening of postmortem kidney confirmed 14 of the enrolled sea otters as *Leptospira-*positive, and the remaining five were negative. Based on cumulative results from necropsy, as well as renal and hepatic histologic examination and IHC, seven of the 14 PCR-positive sea otters were classified as fatal leptospirosis. In all seven cases *Leptospira*-associated lesions were considered a major cause of stranding or mortality based on lesion severity. The remaining seven PCR-positive SSO were classified nonfatal leptospirosis cases based on non-leptospiral related renal and hepatic changes, negative renal and hepatic IHC, as well as other major causes of stranding and mortality unrelated to leptospirosis. Six of the 14 *Leptospira* PCR-positive SSO (43%) had been previously tagged and monitored as part of ongoing research. Five SSO were classified *Leptospira*-negative cases based on negative *Leptospira* renal PCR, non-leptospiral related histologic changes, and negative IHC.

Most *Leptospira* PCR-positive SSO (79%:11/14) were adults or aged adults while four of five *Leptospira*-negative cases were also in this age class (Table 1). Ten of the 14 (71%) fatal or nonfatal leptospirosis cases were male, whereas three of five (60%) negative cases were males. *Leptospira* PCR-positive cases were distributed throughout the study period (2005-2025). Most (86%; 12/14) PCR-positive SSO stranded from January through June, and of these, the majority (71%, 10/14) stranded during and immediately following the period of peak seasonal precipitation in central California (October through April). Two of the five (40%) negative cases stranded within the same period. Nearly all cases stranded within Monterey Bay and within 5km of estuarine or embayment habitat (17/19), including all fatal leptospirosis cases (Figure 1).

**Figure 1.**
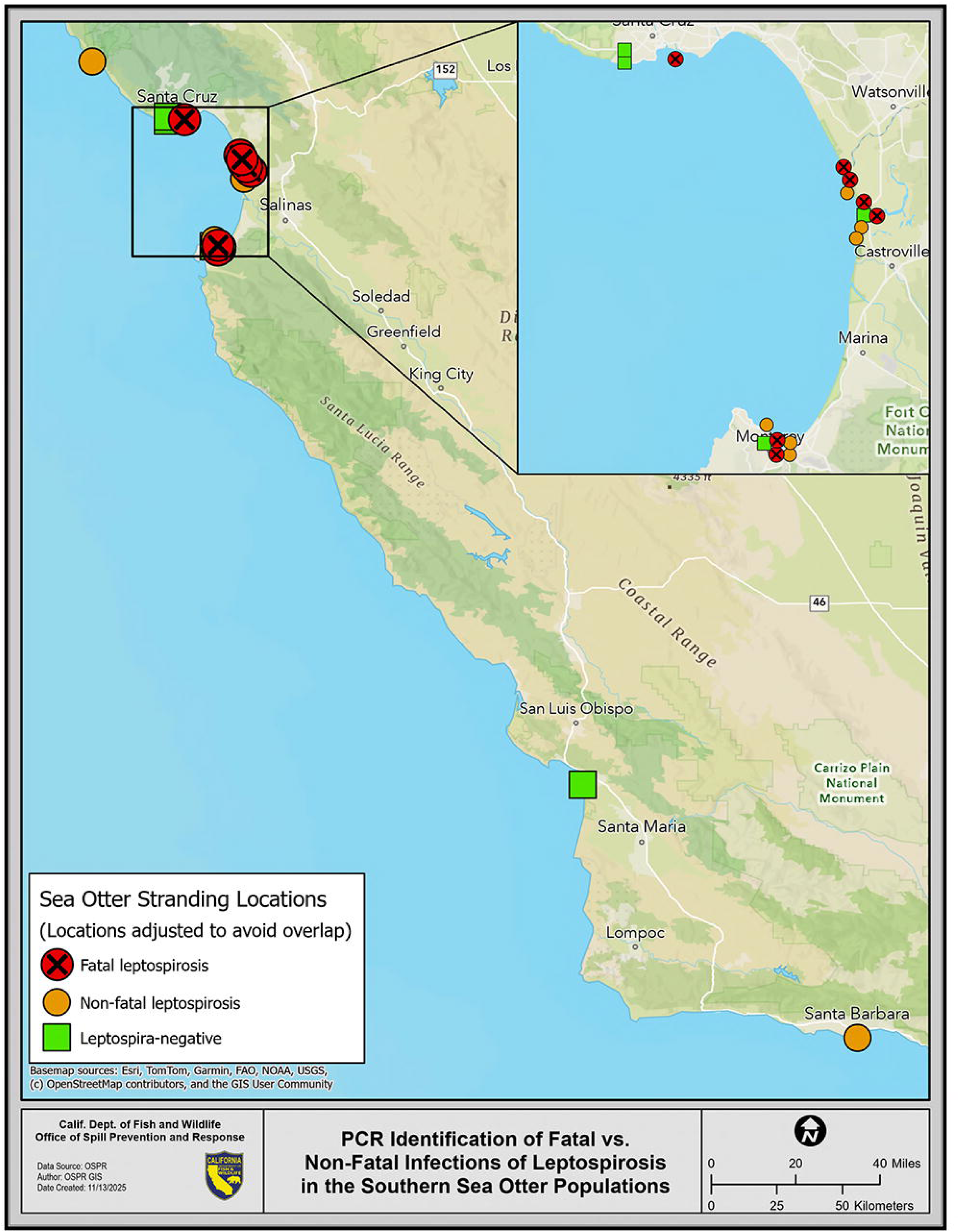
Stranding locations along the central Californian coast for southern sea otters (SSO) that were screened for *Leptospira* infection. **(A)** Distribution of enrolled *Leptospira* PCR-positive and PCR-negative SSO. **(B)** Higher magnification of Monterey Bay Area and distribution of SSO cases with locations adjusted to avoid overlap. Red circle with X = Fatal leptospirosis case. Orange circle = Nonfatal leptospirosis case. Green square = Leptospira-negative case.

### 3.3 Gross, Histologic, and *Leptospira* Immunohistochemical Findings

#### 3.3.1 Fatal leptospirosis cases (N=7)

Of the seven fatal leptospirosis cases, four had grossly abnormal kidneys characterized by randomly distributed miliary white cortical foci throughout all renicules with dark red or light tan cortices (Figure 2A). The remaining three fatal cases had grossly unremarkable kidneys (Figure 2B). Gross hepatic changes were consistent with passive congestion from cardiomyopathy (n=2), malnutrition (n=2) or were unremarkable (Figure 2C). Four of the seven fatal cases had gastric ulcers, a common finding in this population (Miller et al., 2020b), three had oral ulcers and one had concurrent fibrinous posthitis (Table 2). One case was pregnant with a fetus that had grossly apparent congenital malformations, the association with leptospirosis is unknown. Common co-morbidities included malnutrition and cardiomyopathy.

**Figure 2.**
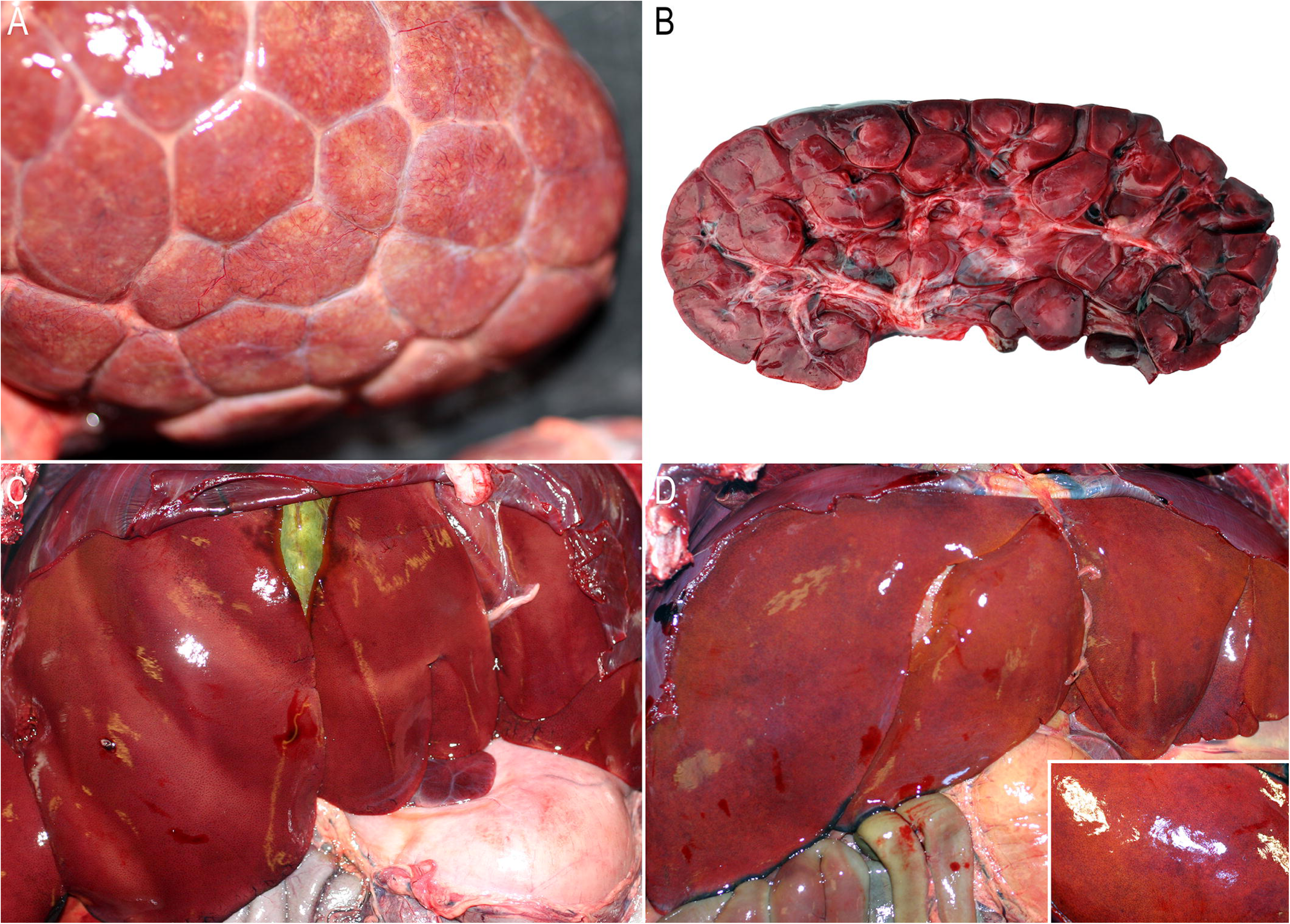
Gross necropsy findings for southern sea otters (SSO) with fatal and non-fatal *Leptospira* infections and PCR-negative animals. **(A)** Kidney from SSO with fatal leptospirosis (case 3) exhibited miliary white subcapsular cortical foci and moderate surface flattening of renicules. **(B)** Grossly unremarkable kidney from SSO with fatal leptospirosis (case 4). **(C)** Grossly unremarkable liver from SSO with fatal leptospirosis (case 2). **(D)** Enlarged friable liver with rounded edges and enhanced reticular pattern (inset) from SSO with fatal microcystin intoxication and nonfatal leptospirosis (case 10). Photo Credits: MWVCRC and TMMC Pathology.

**Table 2.**
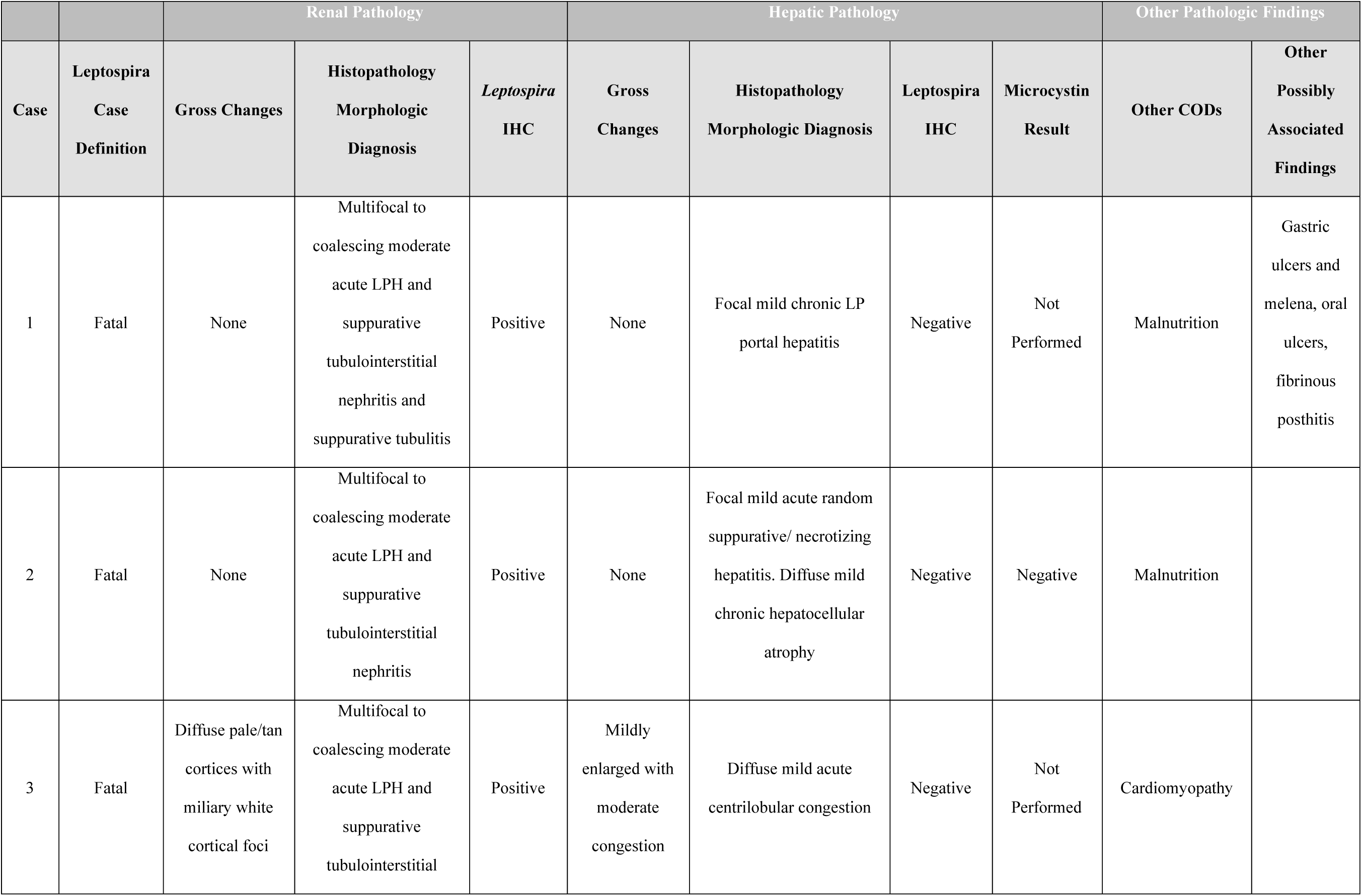
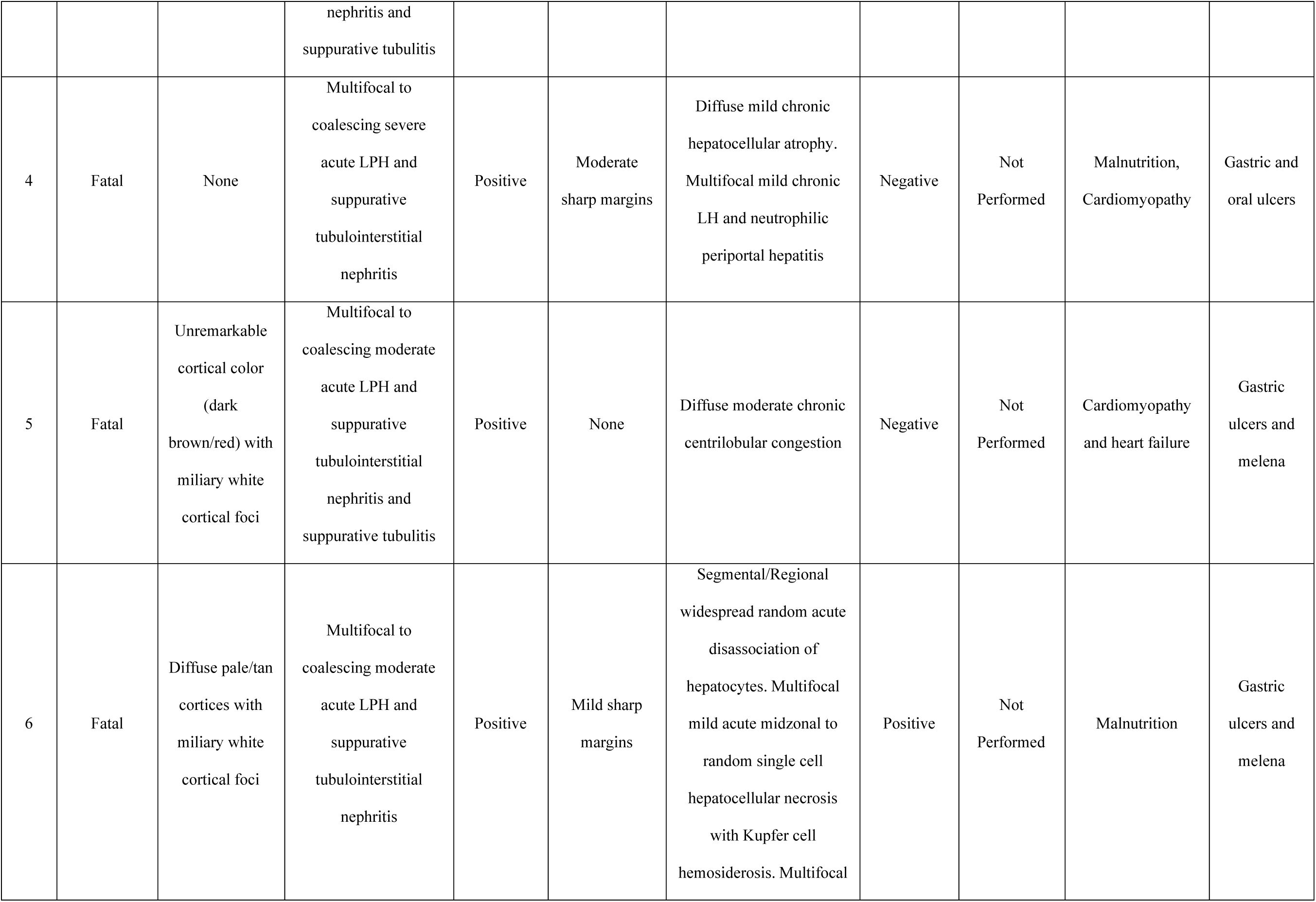
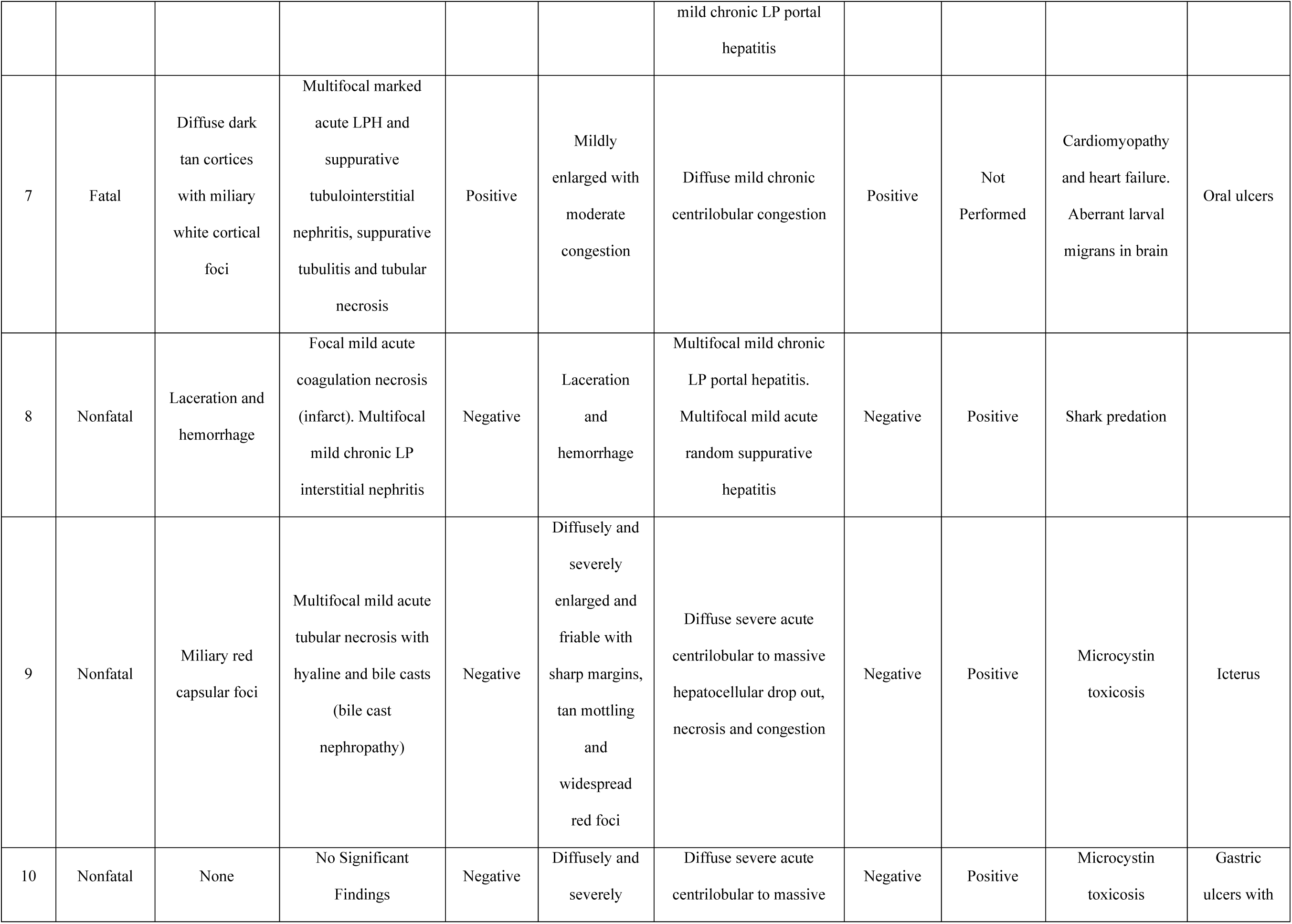
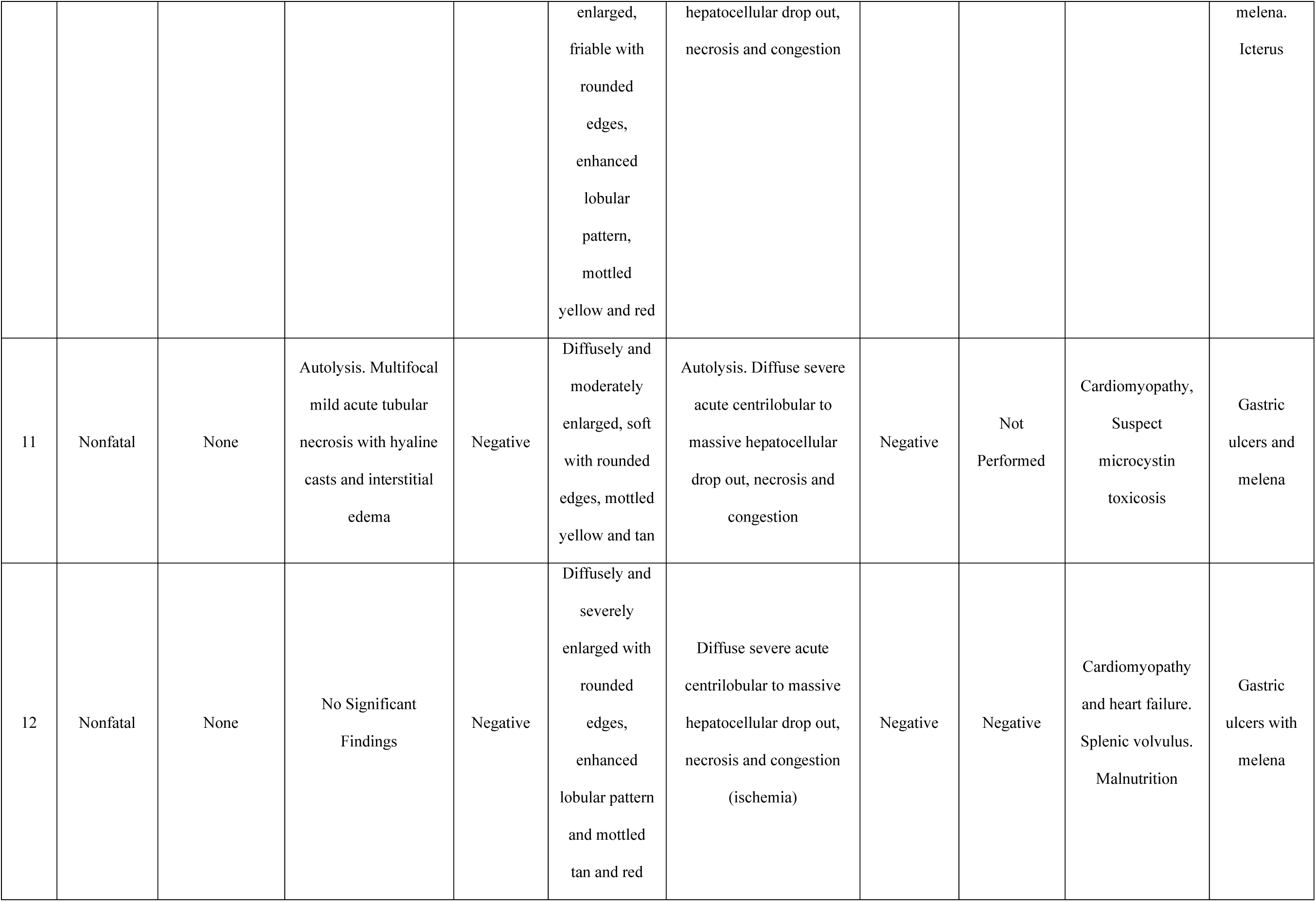
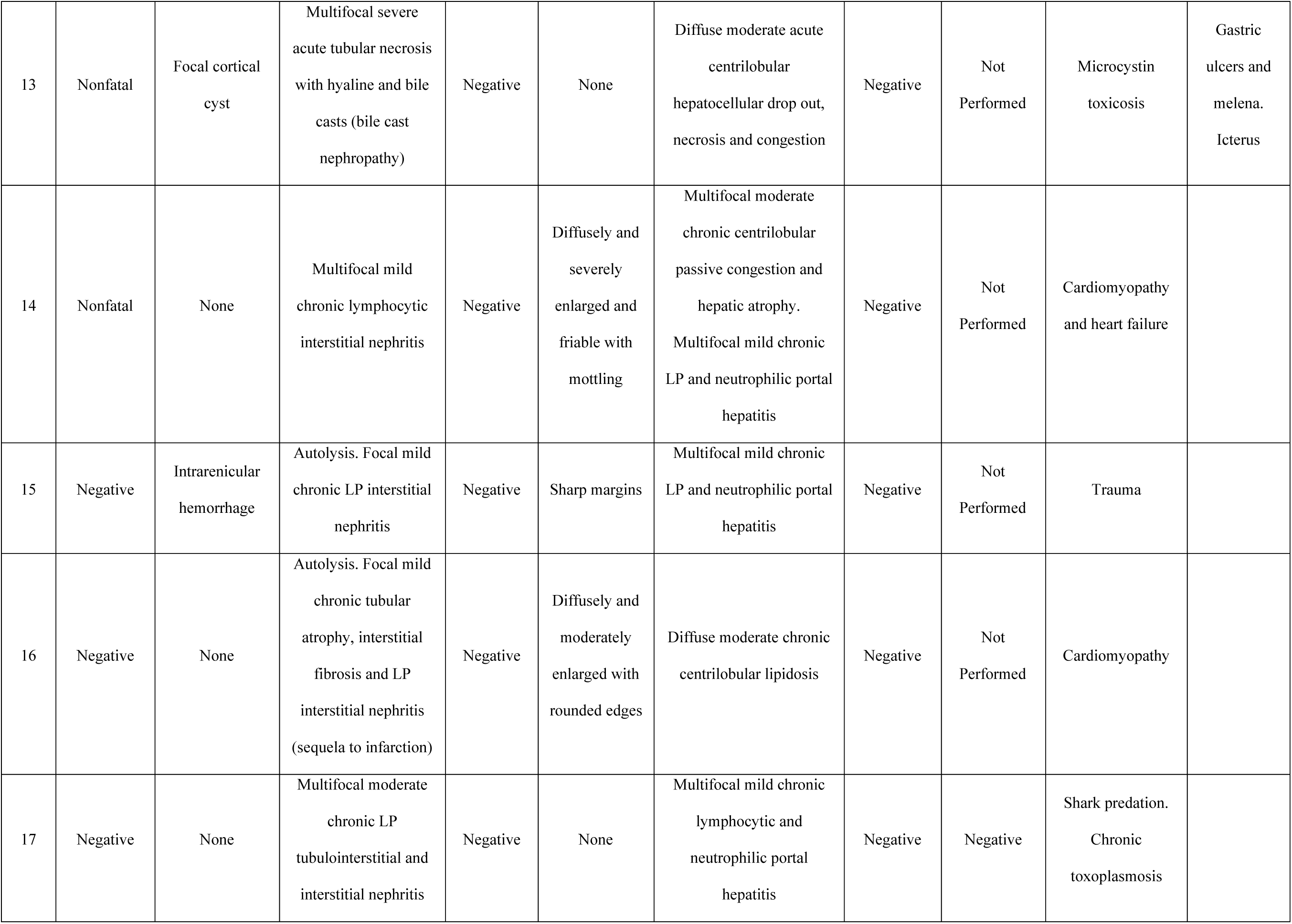
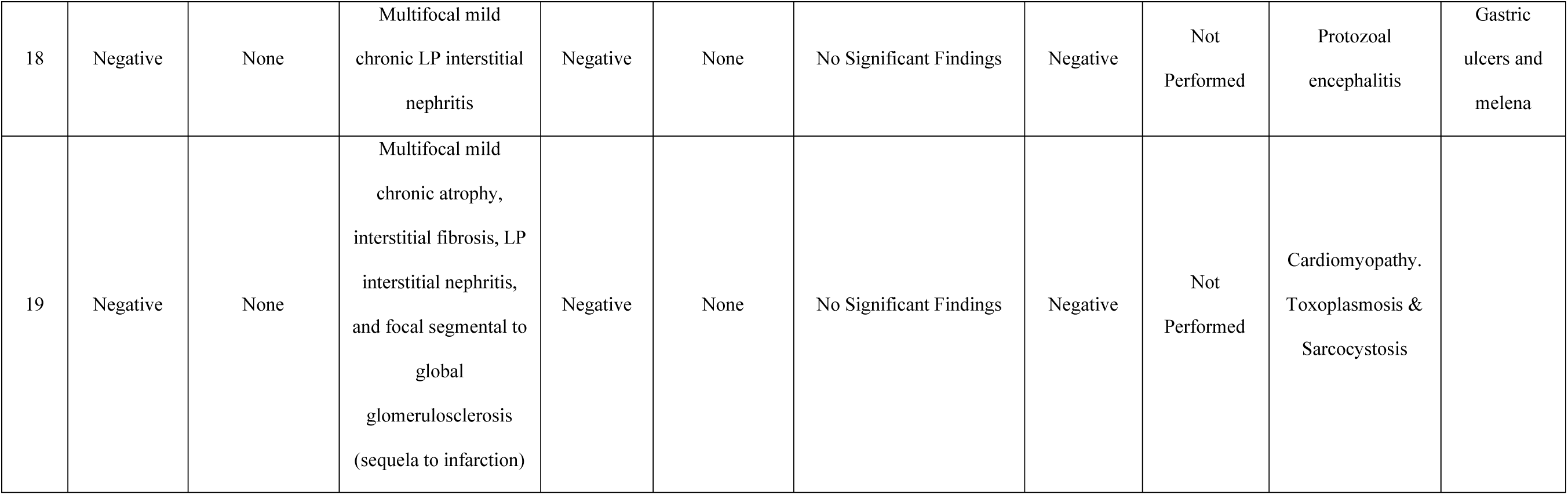
Summary of gross and histopathology findings for fatal and nonfatal leptospirosis and negative southern sea otter cases. LP = lymphoplasmacytic, LPH = lymphoplasmacytic and histiocytic, COD = cause of death.

Histologically, all seven leptospirosis cases with fatal infection had acute moderate to severe lymphoplasmacytic, histiocytic and suppurative tubulointerstitial nephritis with suppurative tubulitis (Figure 3), and positive immunolabeling for *Leptospira* spp.. Tubulointerstitial nephritis was comprised of lymphocytes, plasma cells, histiocytes, and neutrophils expanding the interstitium and dissecting between tubular epithelial cells, with variable rupture of the tubular basement membrane. In some areas renal tubules were completely effaced by inflammatory cells. Most often the tubulointerstitial nephritis was comprised of lymphocytes, plasma cells and histiocytes in the cortex, and neutrophils in the outer medulla.

**Figure 3.**
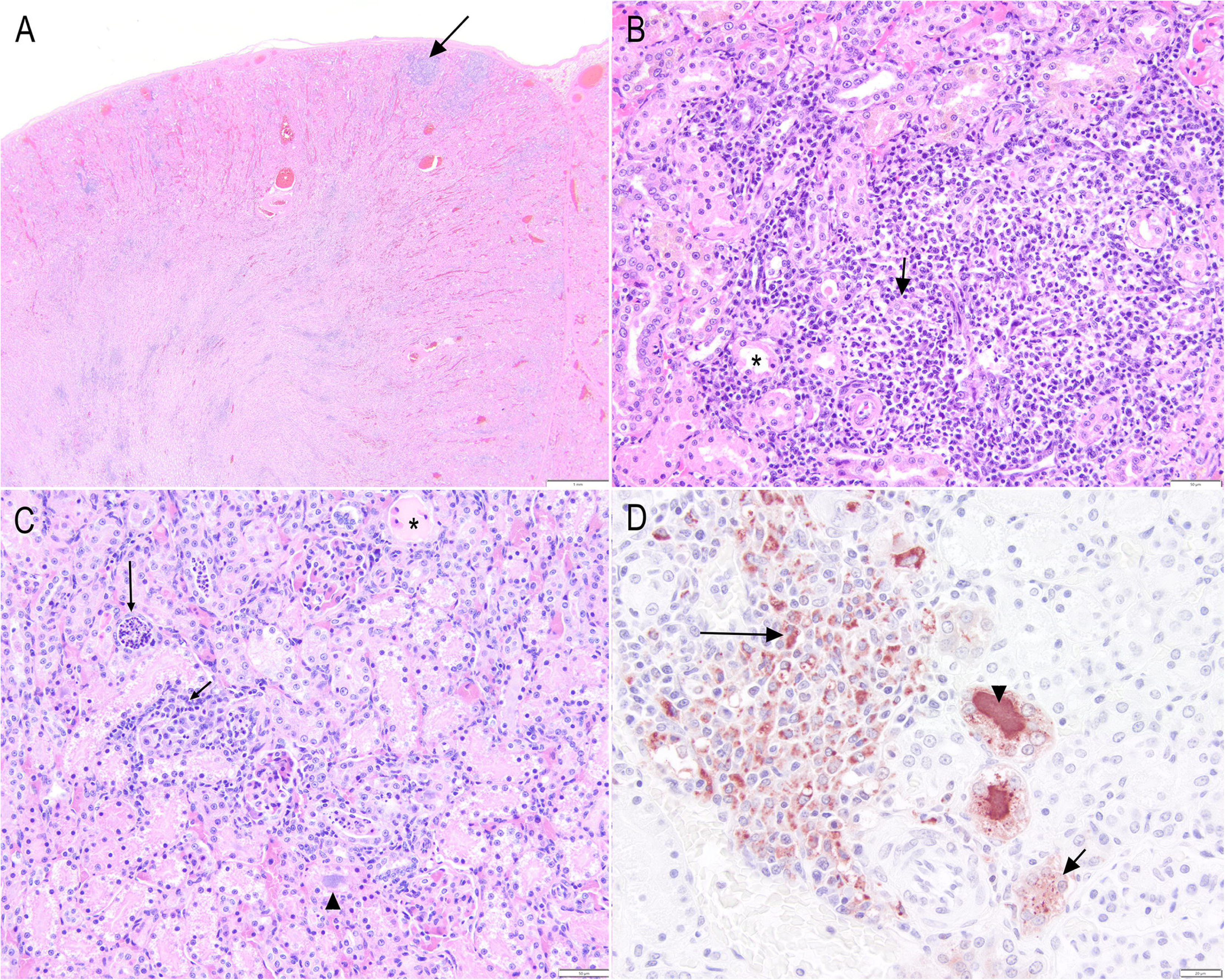
Hematoxylin and eosin-stained tissue sections (H&E) and immunohistochemistry (IHC) from fatal southern sea otter leptospirosis cases. **(A)** Kidney (case 3): Multifocal inflammatory aggregates (arrow) correspond with miliary white subcapsular cortical foci at gross necropsy. **(B)** Higher magnification of a subcapsular inflammatory focus (case 3) showing tubular basement membrane rupture, tubular effacement associated with infiltration by lymphocytes and plasma cells (arrow), and acute tubular necrosis and attenuated epithelium (asterisk). **(C)**. Renal cortex (case 4) characterized by acute tubular necrosis (asterisk), tubulitis with intraluminal cellular debris and degenerate inflammatory cells (long arrow), and tubulointerstitial nephritis comprised of lymphocytes, plasma cells and neutrophils that infiltrate from the interstitium across the tubular basement membrane, dissecting between tubular epithelial cells (short arrow). Tubular lumens are frequently filled with bacteria (arrowhead). **(D)** IHC of renal cortex (case 4) labeling leptospiral organisms with strong granular brown pigment within the cytoplasm of inflammatory cells (long arrow), tubular lumens (arrowhead), and punctate foci within the cytoplasm of tubular epithelium (short arrow). Photo Credits: TMMC Pathology.

The tubulointerstitial nephritis was often most severe in the outer medulla, with the cortex and corticomedullary junction also frequently affected. Tubulitis was characterized by tubular luminal degenerate neutrophils, with or without necrotic sloughed tubular epithelium, and intact tubular basement membranes with variable tubular degeneration, necrosis and attenuation. When present, acute tubular necrosis was characterized by attenuated tubular epithelium with intraluminal sloughed necrotic epithelium. Tubular casts were hyaline or granular, and lumina were often filled with dense colonies of basophilic bacteria. Intraluminal bacteria were common in the papilla, with increased frequency and number associated with progression down the nephron. Other changes included multifocal suppurative vasculitis and fibrin thrombi, especially at the corticomedullary junctions and outer medulla.

*Leptospira* immunolabeling was most often in the cytoplasm of tubular epithelium, specifically the cortical tubular epithelium; however, tubular luminal immunolabeling was mostly within the papilla and corresponded to the bacterial colonies visible in Hematoxylin and eosin-stained (H&E) sections. Inflammatory cells, most likely macrophages, also had positive immunolabeling within foci of tubulointerstitial nephritis, though the tubular and intraluminal staining was more prominent.

Several (3/7) SSO with fatal leptospirosis had mild lymphoplasmacytic, histiocytic and neutrophilic portal to periportal hepatitis that was incidental and unrelated to leptospirosis. One case had a focus of random necrotizing hepatitis that was attributed to septicemia as no leptospiral antigen was present in the focus, while several had passive centrilobular congestion presumed secondary to cardiomyopathy (3/7) or hepatocellular atrophy secondary to malnutrition (2/7); however, one case with fatal leptospirosis had disassociation of hepatocytes with single cell necrosis that had rare, intense granular cytoplasmic staining of individual cells suspected to be individualized hepatocytes or Kupffer cells. Another fatal SSO had rare moderate to intense granular cytoplasmic staining of individual cells, suspected to be Kupffer cells, without any concurrent hepatic changes. All other livers (5/7) had no immunolabeling for *Leptospira* organisms. One fatal leptospirosis case had thrombosis of gastric submucosal vasculature that was associated with mucosal ulceration.

#### 3.3.2 Nonfatal leptospirosis cases (N=7)

Four of the nonfatal *Leptospira*-infected SSO had grossly unremarkable kidneys, while three had unrelated incidental gross findings (Table 2). One sea otter with nonfatal *Leptospira* infection had gross changes consistent with trauma, four had friable, enlarged and mottled livers consistent with microcystin toxicosis (Figure 2D), and two had enlarged congested livers. Four of the seven cases had gastric ulcers, but none had oral ulcers, and three had icterus presumably related to concurrent microcystin intoxication rather than leptospirosis.

Histologically, two of the nonfatal *Leptospira*-infected cases had mild non-specific, incidental lymphoplasmacytic interstitial nephritis, one of which also had an acute focal infarct attributed to hypoxia associated with acute blood loss from trauma. Three cases had bile cast nephropathy related to microcystin toxicosis, while two had no substantial microscopic changes. In the liver, two cases had incidental mild lymphoplasmacytic and neutrophilic portal hepatitis, one of which also had mild to moderate random suppurative hepatitis attributed to septicemia.

Four cases had centrilobular to massive necrosis and hemorrhage and/or congestion consistent with microcystin toxicosis; three of which had biochemical testing for microcystin performed and were positive. One case had centrilobular necrosis attributed to ischemia related to splenic volvulus and tested negative microcystin, and another with centrilobular congestion consistent with passive congestion secondary to cardiomyopathy. None of the seven SSO with nonfatal *Leptospira* infections detected via renal PCR had immunolabeling for leptospiral antigen in kidney or liver.

#### 3.3.3 *Leptospira-*negative cases (N=5)

Only one *Leptospira* PCR-negative SSO had gross renal changes, which were attributed to trauma. An additional negative case had hepatic changes consistent with malnutrition, while a third had hepatic changes consistent with sudden negative energy balance. One of the five negative cases also had gastric ulcers, and none had oral ulcers. All five cases had incidental mild to moderate lymphoplasmacytic interstitial nephritis that did not have a strong component of tubular inflammation and effacement. Two cases also exhibited chronic tubular loss and interstitial fibrosis consistent with sequela to previous renal infarction. In the liver, two cases had incidental lymphoplasmacytic, histiocytic and neutrophilic portal hepatitis, one with moderate lipidosis and the other two with no other findings. None of the five *Leptospira* PCR-negative SSO had positive immunolabeling for leptospiral organisms in the kidney or liver.

### 3.4 Clinical Presentation, Imaging and Hematologic Values

#### 3.4.1 Fatal leptospirosis cases

One fatal case had antemortem renal ultrasound performed (Figure 4) on the day of stranding which was two days prior to death and necropsy. Ultrasonographic imaging revealed an unremarkable urinary bladder wall with echogenic urine, enhanced corticomedullary distinction in the kidney and diffusely increased echogenicity of the cortices with mildly dilated pelvises and calyces (Fig 4B). This animal also presented with severe oral ulcers (Fig 4C).

**Figure 4.**
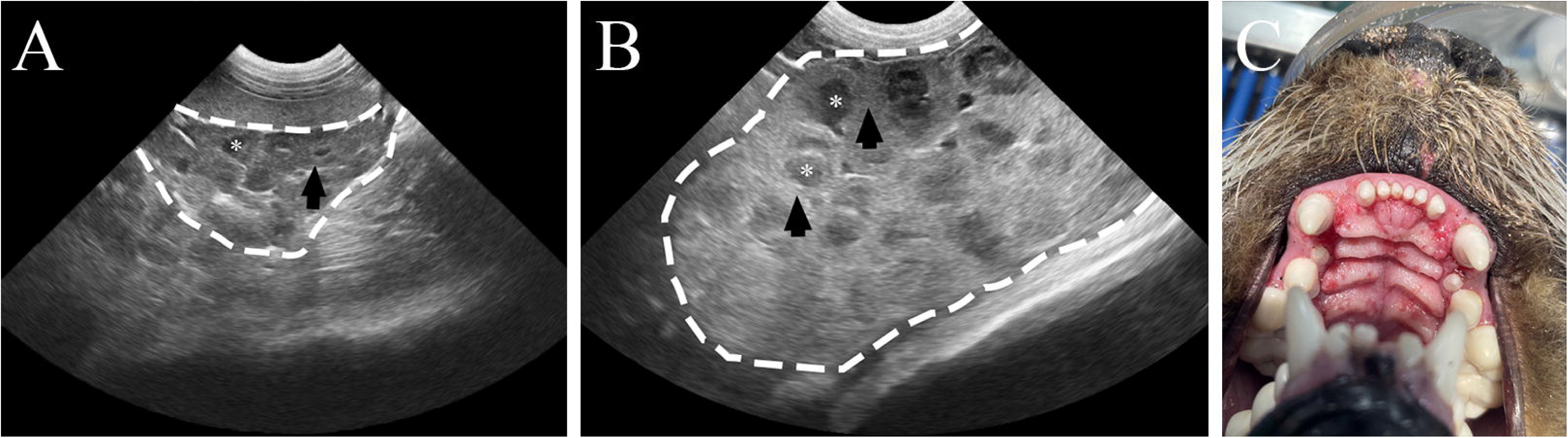
Physical examination and renal ultrasonographic (U/S) findings of a southern sea otter (SSO) with fatal leptospirosis (case 7) compared to a live SSO under human care with unremarkable bloodwork and no evidence of kidney disease. **(A)** Unremarkable renal U/S from clinically healthy SSO. **(B)** Antemortem renal U/S of case 7 with fatal leptospirosis with increased corticomedullary distinction due to hyperechoic cortices. **(C)** Oral ulcerations attributed to uremia at physical examination of SSO with fatal leptospirosis (case 7). Dashed line = outline of renal capsule. Arrow = cortex. Asterisk = medulla. Photo Credits: MBA Veterinary Team.

Five of the seven fatal leptospirosis cases had antemortem bloodwork characterized by mild to moderate azotemia, hypernatremia, hyperkalemia, and hyperphosphatemia (Table 3) (Williams and Pulley, 1983). A complete blood count for case 4 exhibited mild leukocytosis with mild toxic change and left shift (data not shown). Cases 1 and 4 had elevated aspartate aminotransferase (AST), which did not correlate with any hepatic histologic changes and was attributed to skeletal muscle injury during stranding. Both cases also had moderately or markedly elevated creatine kinase in both cases (data not shown).

**Table 3.**
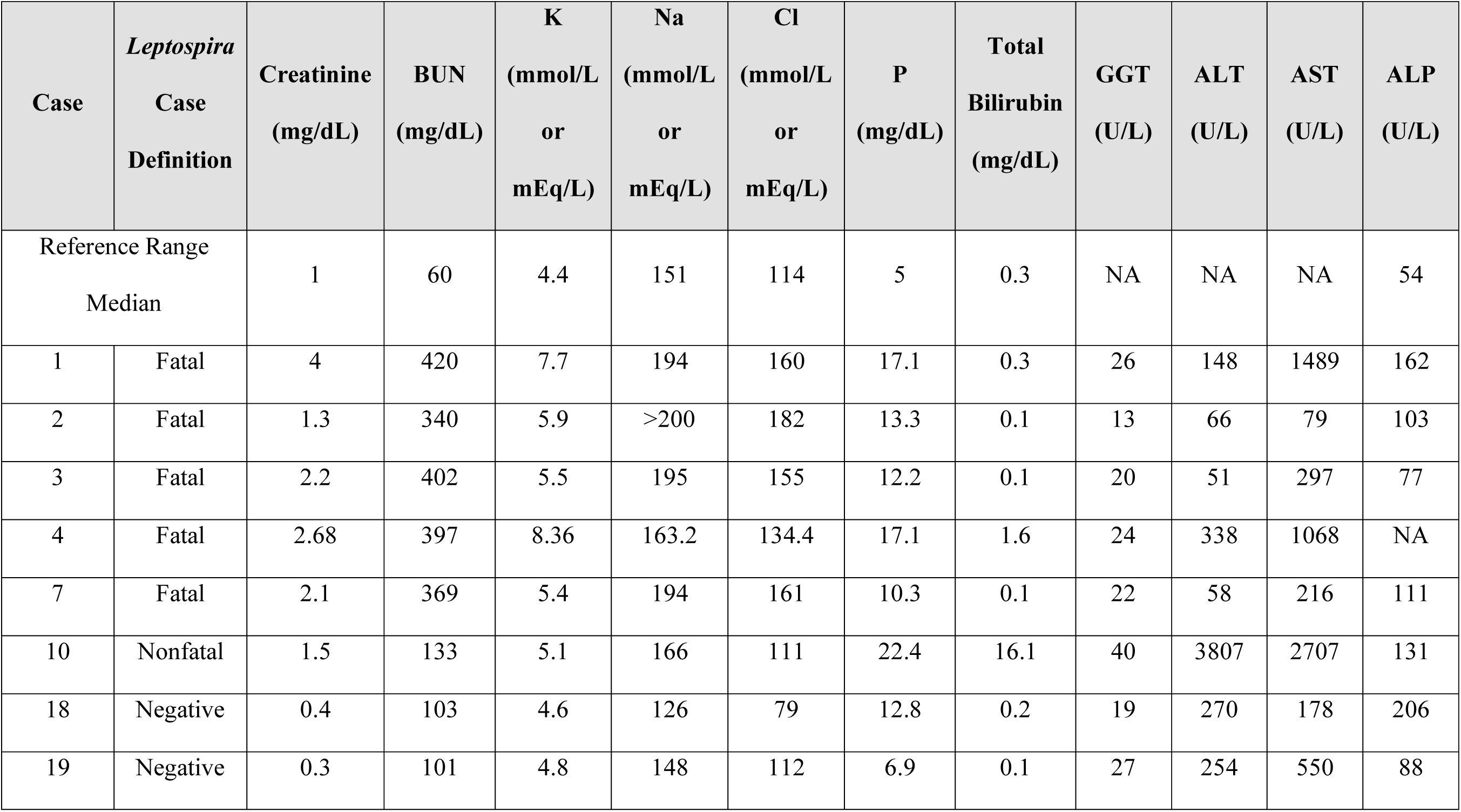
Serum biochemistry values evaluating renal and hepatic function in live-stranded, *Leptospira-* infected and uninfected southern sea otters. Reference range medians from (Williams and Pulley, 1983). BUN= blood urea nitrogen, K= potassium, Na= sodium, Cl= chloride, P = phosphorous, GGT= gamma-glutamyl transferase, ALT= alanine transaminase, AST= aspartate aminotransferase, ALP= alkaline phosphatase, NA= not available.

#### 3.4.2 Nonfatal leptospirosis cases

Case 10, had mild azotemia with hypernatremia, hyperkalemia, marked total hyperbilirubinemia, and elevated ALT and AST. Renal histologic examination did not explain the azotemia and electrolyte abnormalities, which were attributed to dehydration and secondary submicroscopic renal effects from fatal microcystin toxicosis.

#### 3.4.3 *Leptospira-*negative cases

Two *Leptospira*-negative cases had unremarkable renal and hepatic parameters. Both had moderately elevated BUN without concurrent elevated serum creatinine, attributed to stress-associated perimortem gastric ulceration and/or dehydration.

### 3.5 *Leptospira* Serologic Values

Microscopic agglutination tests (MAT) for multiple leptospiral serovars were performed for all enrolled SSO with minimally to mildly hemolyzed serum (14/19). All seven SSO with fatal leptospirosis had titers ≥1:25,600 against *L. interrogans* serovar Pomona (Table 1). As is typical for this assay, all were also reactive to other serovars, to varying and lower titer levels (Lloyd-Smith et al., 2007; Mummah et al., 2024). Of the seven non-fatal cases, three had available non-hemolyzed serum. One was seronegative, and two were seropositive at low titers. Case 13 was seropositive to Pomona at 1:400, but also to *L. interrogans* serovar Icterohaemorrhagiae (1:3,200) and serovar Bratislava (1:800); case 12 was seronegative to Pomona and weakly positive to Icterohaemorrhagiae (1:100). Of the four PCR-negative SSO where serum testing was possible, two had low positive titers for *L. interrogans* serovar Icterohaemorrhagiae (1:100) and Hardjo (1:200) and the remaining two were negative.

### 3.6 Leptospiral PCR and typing

PCR results were used to determine infection status in combination with results of histologic examination and immunolabeling. Therefore, by definition, leptospiral DNA was detected in all seven cases with fatal leptospirosis, and all seven with nonfatal *Leptospira* infections, and in none of the five non-*Leptospira* cases (Table 1). Eight of the 14 *Leptospira* PCR-positive cases, which included six fatal and two nonfatal cases, had DNA sequence-based serogroup typing performed that determined *L. interrogans* serogroup Pomona as the serogroup for all eight cases (GenBank accession numbers: BankIt3042564 Seq1 PX925413, BankIt3047383 Seq1 PX925414, BankIt3047384 Seq1 PX925415, BankIt3047386 Seq1 PX925416, BankIt3047390 Seq1 PX925417, BankIt3047391 Seq1 PX925418, BankIt3047392 Seq1 PX925419, BankIt3047394 Seq1 PX925420). The remaining six of the 14 *Leptospira* PCR-positive cases did not yield good quality DNA sequences, precluding serogroup identification.

## 4 Discussion

This is the first report of *Leptospira* spp. infection and fatal leptospirosis in SSOs from California. Although there are similarities with prior reports of leptospirosis in marine mammals, terrestrial mammals, and humans, with respect to the associated gross and microscopic lesions, there are also some unique differences.

As with CSLs, SSOs infected with *Leptospira* spp. were more often males (Emily R Whitmer et al., 2021), and similar findings were reported for NSOs with leptospirosis from Washington state (Knowles et al. 2020). Whether a true sex predilection and the reason for it in SSO is not known and additional epidemiologic studies could provide definitive insights. There is little segregation by sex in SSOs, with males tending to concentrate in shallow areas with sandy embayments, such as Monterey Bay and Estero Bay, and females concentrated in nearshore kelp covered areas (Jameson, 1989; Tinker et al., 2017). Moreover, males are known to range more widely than females (Ralls et al., 1996; Tinker et al., 2008); as a result, they could be more likely to encounter coastal point sources for *Leptospira* exposure. In contrast, there is marked sexual segregation among CSLs along the California coast outside of the summer breeding season and leptospirosis outbreaks have been dominated by infections in males in central and northern California (Gulland et al., 1996).

The lesions of *Leptospira* infection in SSO was characterized by a mix of similarities and differences compared to other host species. Southern sea otters with fatal leptospirosis exhibited unique and highly variable gross renal changes; some cases had miliary white cortical foci, and other cases had grossly unremarkable kidneys. Northern sea otters with fatal leptospirosis also had subtle non-specific gross renal changes, such as swelling and congestion, but pale streaks were also reported that could be consistent with the white miliary subcapsular foci observed in the present study (Knowles et al., 2020). All gross features of SSO leptospirosis were distinct from the striking diffuse renomegaly and cortical pallor characteristic of leptospirosis in CSLs (Gulland et al., 1996). Therefore, diagnosis of leptospirosis in SSO should not be based solely on gross appearance of the kidneys at necropsy.

SSO with fatal leptospirosis primarily had histologic lesions characterized by tubulointerstitial nephritis with intralesional leptospiral organisms confirmed via IHC. These histologic features are also characteristic of leptospirosis in NSOs, CSLs, northern elephant seals (NES, *Mirounga angustirostris*), and harbor seals (HS, *Phoca vitulina*) (Delaney et al., 2014; Gulland et al., 1996; Knowles et al., 2020; Stamper et al., 1998a). In addition, for SSO cases reported here, the tubulointerstitial nephritis was distributed throughout the renal cortex, corticomedullary junction, medulla and papilla, but was most prominent in the outer medulla. By contrast, leptospirosis is traditionally described in domestic host species as cortical and perivascular corticomedullary tubulointerstitial nephritis with the predominant infiltrates being lymphocytes and plasma cells (Maxie, 2016). However, common concurrent features of leptospirosis in our study included suppurative tubulitis and acute tubular necrosis, which have been infrequently described in other marine mammals but are also commonly observed in pinniped leptospirosis at TMMC (Martinez pers. observation). One potential differential for tubulitis in the medulla and papilla is secondary ascending non-leptospiral bacterial urinary tract infection, especially given the prominent intraluminal bacterial colonies seen with H&E. Anecdotally, fatal leptospirosis CSLs at TMMC have also been documented to have concurrent UTI via urine culture and histologic examination (Martinez pers. observation). Aerobic and anaerobic urine cultures were not routinely performed in the otters; therefore, concurrent urinary tract infections cannot be fully ruled out. However, intense positive immunolabeling of tubular luminal contents that coincide with the bacterial colonies on H&E as well as the foci of medullary and papillary tubulitis and tubulointerstitial nephritis suggest that these lesions may be due to proliferating leptospiral organisms. Observable bacterial colonies on H&E are not a commonly described feature in leptospirosis of other species, and this finding could reflect the acute shedding phase of infection in SSOs and/or the relative pathogenicity of certain *Leptospira* strains in sea otters. It may be that SSOs strand earlier in the disease process than CSLs due to their comparatively high metabolic rate and limited energy reserves, or these histologic and IHC findings may indicate particularly intense shedding.

A common incidental and nonspecific finding among nonfatal leptospirosis and negative SSO cases was mild to moderate lymphoplasmacytic interstitial nephritis. This change was associated with little to no tubular damage, negative or very low *Leptospira* titers, and should not be confused with fatal leptospirosis. A differential could be a resolving inflammatory process such as historical leptospirosis; however, given the negative or very low (1:400) *Leptospira* serovar Pomona antibody levels in these cases, and absence of tubular involvement, this is less likely the scenario for these cases. Therefore, as proposed in domestic species, other sources of previous injury, e.g. regional ischemia, could be the cause of these non-specific foci of interstitial nephritis.

In SSOs, leptospirosis appears to primarily target the kidneys, with infrequent manifestation of hemodynamic disturbances or hepatic disease. Anecdotally, some of the PCR-positive sea otters, as well as PCR-negative cases, had petechiae in various organs but this was attributed to postmortem changes rather than vasculitis and disseminated intravascular coagulopathy as occurs with leptospiral petechiation in other species and following infection with serovars other than Pomona (Haake and Levett, 2015; Levett, 2001). However, one fatal case did have thrombosis that was associated with gastric ulceration and may represent *Leptospira*-associated lesion. When petechiae were present in animals from this case series, it was most often associated with microcystin toxicosis rather than *Leptospira* infection. Similarly, icterus, which can be associated with leptospirosis in other species (Levett, 2001), was associated here with microcystin toxicosis in SSOs (Levett, 2001). Gastric ulceration was commonly observed in fatal and nonfatal leptospirosis cases. Although this condition could be due to uremia secondary to leptospirosis, or thrombosis associated with leptospirosis as in one fatal case, gastric ulceration is a common finding in stranded SSO and is presumed to be due to stress (Miller et al., 2020b). In contrast, ulcers of the oral mucosa were only observed in three fatal leptospirosis cases and are otherwise uncommon in stranded SSOs. Oral ulceration was not present in nonfatal leptospirosis cases nor *Leptospira-*negative cases, suggesting that this lesion could indicate uremia due to renal failure as it does in CSL (Duignan et al., 2019).

In other host species, leptospirosis has been associated with hepatic disease that ranges from hepatitis to disassociation of hepatocytes related to a direct effect of leptospires on intercellular tight junctions (Maxie, 2016). One SSO with fatal leptospirosis exhibited disassociation of hepatocytes with rare positive immunolabeling of leptospires. This classic hepatic lesion in terrestrial animals has not been previously reported in marine mammals with leptospirosis and appears to be a rare finding in SSO (Delaney et al., 2014; Gulland et al., 1996; Haake & Levett, 2015; Rajapakse, 2022; Samrot et al., 2021; Stamper et al., 1998a). A few *Leptospira*-infected SSOs exhibited marked centrilobular coagulation necrosis and congestion associated with biochemically confirmed microcystin toxicosis; this lesion could be confused with hepatic lesions of leptospirosis and warrants further investigation when such lesions are detected. The subtle change of disassociation of hepatocytes could be confused with early or mild microcystin intoxication, edema or autolysis; however, when present, the hepatic lesion associated with microcystin toxicosis was severe centrilobular to massive necrosis and hemorrhage. Further differentiation of microcystin toxicosis from leptospirosis may be achieved by concurrent evaluation of the renal and hepatic changes, such as the presence of tubulointerstitial nephritis, which would support leptospirosis, while acute tubular necrosis and/or bile cast nephropathy would support microcystin toxicosis (Xu et al., 2020). It is also noteworthy that three SSOs had concurrent fatal microcystin toxicosis and incidental *Leptospira* infection, so it may be best to screen for both conditions in challenging cases. Lastly, it is worth noting that the severity of hepatic lesions due to microcystin toxicosis could have obscured any subtle changes, such as disassociation of hepatocytes related to leptospirosis. With the relatively common concurrent nonfatal leptospirosis infection and microcystin toxicosis, additional studies may be warranted to determine whether one disease predisposes SSO to the other.

Though hepatic lesions are commonly reported in domestic species with leptospirosis, the changes can be non-specific, subtle and/or background lesions such as portal infiltrates and single cell necrosis (Greenlee et al., 2005). The comparatively rare hepatic lesions in SSOs and other marine mammals with leptospirosis could be due to differences in dominant *Leptospira* serovars circulating in the marine environment compared to those commonly causing more obvious hepatic lesions in domestic species, host-specific differences, or the presence of common co-morbidities complicating the ability to distinguish direct *Leptospira* effects from septicemia or endoparasitism. The majority of *Leptospira*-infected SSOs had hepatic changes attributed to incidental portal inflammatory infiltrates from ascending pathogen stimulation, random hepatitis, or septicemia, but with no positive immunolabeling for leptospiral organisms, suggesting an alternate cause. Many SSOs also had centrilobular congestion and hepatic atrophy consistent with right sided heart failure or malnutrition. These are common findings in free ranging SSOs (Miller et al., 2020) due to various causes such as domoic acid toxicosis and protozoal infections and should not be confused with *Leptospira-*associated hepatic disease.

Though not the focus of the current study, we also summarized available *Leptospira-*associated clinical findings, especially those that correlated with lesions and could facilitate antemortem diagnosis and screening in SSOs. Renal ultrasound performed for one SSO with fatal leptospirosis revealed increased corticomedullary distinction and increased cortical echogenicity, similar to findings in CSLs with leptospirosis (Field et al., 2021). Another similarity was that SSOs presented clinically with azotemia, related electrolyte abnormalities, and oral ulcers, though not as severe as in CSLs (Duignan et al., 2019; Whitmer et al., 2021).

Although gross findings for fatal leptospirosis in SSOs can be subtle, antemortem imaging and bloodwork can greatly facilitate diagnosis, which highlights the importance of perimortem hematological screening whenever possible. Interestingly, of the seven PCR-positive nonfatal leptospirosis SSO cases, only one had bloodwork available, with mild azotemia and no histologic lesions suggesting dehydration as the cause. Whether this case represents a subclinical/asymptomatic infection is not known. Further studies could fully characterize the clinical presentation of leptospirosis in SSOs and validate antemortem diagnostic tests used for CSLs (Field et al. 2021; Peters et al., 2025; Whitmer et al., 2021). Similarly, additional studies could determine the possibility of chronic renal tubular colonization or chronic shedding in the nonfatal leptospirosis cases and the potential of such cases to be reservoirs of infection in the population.

In this study, LigB amplification and sequencing found that all fatal leptospirosis cases were infected with *L. interrogans* serogroup Pomona. This was supported by serological findings, with all fatal cases exhibiting peak titers against serovar Pomona. Taken together, this evidence is consistent with Pomona being the dominant serovar infecting SSOs, similar to CSLs, a population in which it has circulated for decades (Prager et al., 2020). Because of cross reactivity, serologic tests alone cannot confirm the infecting serovar, necessitating genetic sequencing to facilitate population level epidemiologic studies (Mummah et al., 2024). Although there were a few positive titers for other serovars, most were lower than the titer to Pomona and likely represent cross reactivity (Mummah et al., 2024). One intriguing outlier was a nonfatal case (case 13) with a MAT titer of 1:3200 against *L. interrogans* serovar *Icterohemorrhagiae*, compared to 1:400 against serovar Pomona. However, paradoxical MAT reactions (i.e. higher titers against serovars other than the infecting serovar) are sometimes reported, and even an 8-fold difference in titer is unusual but not unprecedented (Mummah et al., 2024). *Leptospira interrogans* serovar Pomona has also caused infections and disease in NESs, HSs and NSOs (Delaney et al., 2014; Knowles et al. 2020; Prager et al., 2013; Stamper et al., 1998a; Zuerner & Alt, 2009, Knowles et al., 2020). Furthermore, this serovar also circulates in coastal mammals including raccoons and coyotes (Helman et al., 2023; Straub et al., 2020; Straub and Foley, 2020).

Although we observed some intriguing temporal clustering of *Leptospira*-infected SSOs, we have interpreted this cautiously because our sample size was small, case inclusion was biased, and the focus of the current study was case confirmation and description of lesions rather than epidemiologic analysis. Most *Leptospira*-positive SSOs stranded during the first half of the year, especially during, or soon after, the winter rainy season. This could reflect exposure via runoff and land-sea transmission, as has been reported for apicomplexan protozoal infections, selected metazoan parasites, and other bacterial species infecting SSOs (de Wit et al., 2020; Miller et al., 2020a, 2020b, 2002; Oates et al., 2012; Shapiro et al., 2012). However, late winter and spring are the peak season for SSO strandings, so additional study would help to confirm seasonal patterns of stranding for *Leptospira-*infected SSO in relation to rainfall and land-sea runoff. It is also possible that *Leptospira* may transmit between CSLs and SSOs, given that both hosts share the same serovar. It is noteworthy that six of the seven fatal leptospirosis SSO cases stranded in years when large outbreaks occurred among sympatric sea lions. Case 4 stranded in a region (Del Monte Beach, Monterey) that was also heavily occupied by CSLs and potential *Leptospira* transmission from CSLs to other sympatric marine mammal species has been suggested (Colegrove et al., 2005; Delaney et al., 2014; Stamper et al., 1998a). However, three of the 14 *Leptospira-*infected SSO cases, including one fatal case, occurred during a four-year period (2013 to mid-2017) when *Leptospira* was absent from the CSL population (Lloyd-Smith et al., 2023), suggesting that transmission to SSOs may not solely be driven by exposure to infected CSLs. Considering that seven SSOs had nonfatal infections, it is also possible that *Leptospira* could be maintained in the SSO population and continuing exposure could be due to chronic or asymptomatic shedders as in CSLs and other mammals (Buhnerkempe et al., 2017; Lloyd-Smith et al., 2007; Prager et al., 2013, 2020). However, these hypothesis-generating observations warrant further investigation.

Apparent spatial clustering was observed, though again the non-random case selection in our study prevents formal analysis or strong conclusions. Nearly all *Leptospira*-infected SSOs stranded within Monterey Bay, though this is the region where most otters stranded that were included in the current pilot study. Similarly, most cases stranded within or adjacent to estuaries, sloughs, embayments, or harbors, though this was also true of enrolled *Leptospira*-negative animals. Both spatial patterns could arise from biased sampling, as these well trafficked areas are more likely to have stranded marine mammals reported by the public to the stranding network.

Nevertheless, there are several attributes of embayment areas that could be considered for additional epidemiologic studies to determine risk factors for efficient transmission of *Leptospira* to and among SSOs. Enclosed water bodies could enhance pathogen exposure through concentration of contaminated freshwater inflows, followed by slow dilution during tidal exchanges. Reduced salinity in these zones could elevate risk for leptospiral infection to SSO as *Leptospira* organisms survive longer in fresh or brackish water (Bierque et al., 2020). Finally, embayments along the California coast provide habitat for multiple marine mammal species living in close proximity. It is noteworthy that prior studies of other pathogens and toxins have reported higher risk for fatal outcomes for SSOs residing in or near certain coastal zones and embayments (Burgess et al., 2020; de Wit et al., 2020; Miller et al., 2010, 2020). These preliminary findings indicate that additional epidemiologic studies could provide clarity concerning the landscapes and spatial risk factors associated with *Leptospira* infection for SSO.

At the time of manuscript preparation, a small-scale outbreak of leptospirosis with multiple fatal cases, including three in the current study, was ongoing in SSO in or near Elkhorn Slough/Moss Landing in central Monterey Bay (Miller pers observation). This outbreak preceded, then was concurrent with, an outbreak of unprecedented magnitude among CSLs along the central California coast. This observation heightens the urgency of untangling the possible transmission links between these sympatric marine mammal species.

In conclusion, this is the first confirmation of *Leptospira* infection and fatal leptospirosis in SSOs from California. Additional studies could clarify the prevalence of *Leptospira* infection and fatal leptospirosis in the population. This study revealed similarities and differences between *Leptospira* associated disease in SSO compared to pinnipeds and terrestrial species and provided insight into characteristics to facilitate case detection in future research and clinical settings.

Although intriguing preliminary spatial and temporal patterns were detected, the present study design was not suitable for formal statistical analysis. Larger and more systematic studies could assess these epidemiologic patterns and identify transmission pathways between various marine and terrestrial mammal populations, and potential anthropogenic drivers of disease incidence and population impacts in this threatened marine mammal.

## 5 Acknowledgements

The authors acknowledge the dedicated volunteers, staff and donors at The Marine Mammal Center, CDFW, and MBA for making this research possible. K.C.P. and J.O.L.-S. were supported by the US National Science Foundation (DEB-1557022. K.C.P. was supported by the John H. Prescott Marine Mammal Rescue Assistance Grant Program (NA23NMF4390341) and US Fish and Wildlife Service American Rescue Plan Act Zoonotic Disease Initiative (F23AP00118-00). The Marine Mammal Center was supported by the USFWS John H. Prescott Marine Mammal Rescue Assistance Grant Program Grant #F24ASOO186 and #F23AP02422. J.T. was supported by the U.S. Geological Survey Ecosystems Mission Area. The authors would like to also thank and acknowledge the work of Francesca Batac, Andrew Caputo, Michelle Staedler, Renee Galloway, CDC, CAHFS, and MSU VDL that contributed to the study. Data either are not available or have limited availability owing to proprietary interest of the funding agency: John H. Prescott Marine Mammal Rescue Assistance Grant Program Grant. Contact Mellissa Miller for more information. Any use of trade, firm, or product names is for descriptive purposes only and does not imply endorsement by the U.S. Government.

